# Variations in visual sensitivity predict motion sickness in virtual reality

**DOI:** 10.1101/488817

**Authors:** Jacqueline M. Fulvio, Mohan Ji, Bas Rokers

## Abstract

Severity of motion sickness varies across individuals. While some experience immediate symptoms, others seem relatively immune. We explored a potential explanation for such individual variability based on cue conflict theory. According to cue conflict theory, sensory signals that lead to mutually incompatible perceptual interpretations will produce physical discomfort. A direct consequence of such theory is that individuals with greater sensitivity to visual (or vestibular) sensory cues should show greater susceptibility, because they would be more likely to detect a conflict. Using virtual reality (VR), we first assessed individual sensitivity to a number of visual cues and subsequently induced moderate levels of motion sickness using stereoscopic movies presented in the VR headset. We found that an observer’s sensitivity to motion parallax cues predicted severity of motion sickness symptoms. We also evaluated evidence for another reported source of variability in motion sickness severity in VR, namely sex, but found little support. We speculate that previously-reported sex differences might have been due to poor personalization of VR displays, which default to male settings and introduce cue conflicts for the majority of females. Our results identify a sensory sensitivity-based predictor of motion sickness, which can be used to personalize VR experiences and mitigate discomfort.

## 1. Introduction

Although the visual system is often studied in relative isolation, it has clear connections to other components of the nervous system, for example in the regulation of diurnal rhythm, arousal and balance. One area where this connection is painfully clear is in the domain of motion sickness. However, there is considerable variation in the susceptibility to motion sickness across individuals, and an account for this variability has been elusive.

Cue conflict theory provides a potential account for motion sickness in virtual environments (see [1] for a review). The theory posits that motion sickness is the result of conflict between sensory signals that are typically in concert [2]. Although cue conflicts may arise within a single sensory modality [3], motion sickness is typically attributed to conflicts between visual and vestibular system cues (e.g., [2], [4]-[8]). From an evolutionary perspective, such conflicts were likely to occur following the ingestion of neurotoxins. Thus, the body’s nausea and vomiting responses, which cause the toxin to be expelled, may have developed as the result of an evolutionary adaptation [9]-[12].

There is some support for a relationship between vestibular function and motion sickness. First, motion sickness does not occur in individuals who lack a vestibular system. Second, those with a dysfunctional vestibular system are particularly susceptible [13]. Finally, sensitivity of the vestibular system to self-motion predicts symptoms of motion sickness in individuals with a functioning vestibular system, although the relationship is often small and context-specific [14]. To our knowledge however, a relationship between *visual* sensitivity and motion sickness has not been established.

To account for individual variability in motion sickness severity, we designed a series of experiments. In our experiments, we tested individual observers’ sensitivity (both males and females) to various cues that signal object motion. We manipulated sensory cues pertaining to object motion in depth based on the general visual equivalence between an observer moving through an environment and objects moving around an observer. Prior work has identified considerable variability in the sensitivity to visual cues that specify object motion. For example, observers exhibit independent sensitivity to interocular velocity differences (IOVD) and changing disparities (CD) [15]. Subsequent to our assessments of visual sensitivity, observers watched video footage designed to induce moderate discomfort in a virtual reality headset. We then tested the relationship between sensitivity to various sensory cues and motion sickness due to video viewing.

To summarize our logic, we examined a potential predictor for motion sickness severity. We reasoned that both the vestibular and visual systems provide estimates of the degree of self-motion. If these estimates tend to be highly accurate, then the system should be more likely to detect mismatches between the estimates. Indeed, we did find that the sensitivity to sensory cues to 3D motion predicted an individual’s susceptibility to motion sickness. In particular, individual sensitivity to motion parallax cues produced by small head movements predicts the severity of motion sickness symptoms. In addition, we found evidence that observers self-regulate discomfort by modulating their head movement over time.

We subsequently explored a potential cause for previously reported sex differences in motion sickness susceptibility in virtual reality (VR). Default VR head-mounted display settings tend to be geared toward the average male. Use of default settings for individuals who deviate from the average will introduce cue conflicts into the visual display, and such deviations are of course much more likely for females. Having tailored the display to the interpupillary distance (IPD) of each individual observer, we did not find differences in motion sickness susceptibility based on sex in our sample of observers (see also [16])

Our results suggest a number of strategies to mitigate motion sickness in VR. These strategies include reducing or eliminating specific sensory cues, reducing an observer’s sensitivity to those cues by perhaps counter-intuitively degrading visual fidelity, and ensuring device settings are personalized to each observer.

## 2. Methods

### 2.1. Observers

108 observers were recruited and gave informed written consent. A total of 103 successfully completed all parts of the study. Failure to complete the experiment was either due to technical issues (n = 3), experimenter error (n = 1), or difficulty seeing the stimuli (n = 1). Data from an additional 8 observers were excluded because they did not achieve performance above chance level in any condition on the psychophysical task - see “3D motion discrimination task” section below. Therefore, data from a total of n = 95 observers were included in the main analyses. The required sample size was based on a previous study that investigated motion sickness propensity in virtual reality [17]. The experiments were approved by the Human Subjects Institutional Review Board at the University of Wisconsin-Madison. Observers received course credit in exchange for their participation.

### 2.2. Display Apparatus

Observers viewed visual stimuli in the Oculus Rift Development Kit 2 (DK2; www.oculusvr.com), a stereoscopic head-mounted virtual reality system (see **Fig. 1**, “Virtual reality headset” panel) with a 14.5 cm low-persistence AMOLED screen (Samsung) embedded in the headset providing a resolution of 1920×1080 pixels (960×1080 pixels per eye) with a refresh rate of 75 Hz. The horizontal field of view of the device is about 85 deg (100 deg diagonal). The device utilizes six degrees of freedom (6 DoF) head-tracking for head-motion contingent updating of the display. Positional tracking was achieved at 60 Hz with .05 mm precision via an external camera with a near-infrared CMOS sensor. Tracking of head rotation was achieved at 1000 Hz with .05 deg precision via an accelerometer, gyroscope, and magnetometer embedded in the headset. The effective tracking latency after sensor fusion was about 2 ms and head-movement-to-photon latency was about 14 ms.

**Figure. 1.**
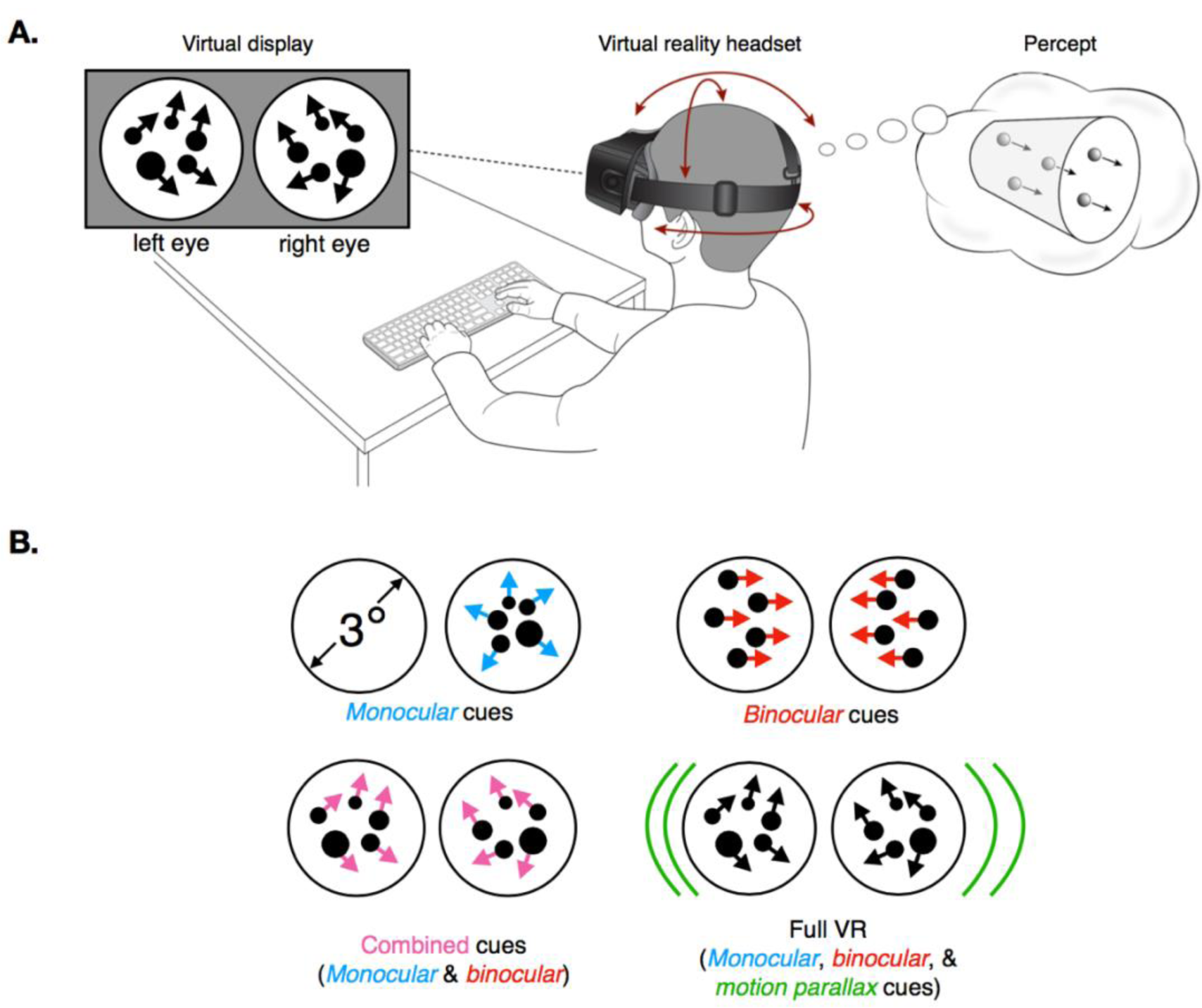
Experimental details. **A**. “Visual display”: Illustration of left- and right-eye stimulus elements as presented in the motion task. The illustration depicts the random dot stimulus. In the actual experiment, the dot stimulus was comprised of 12 dots whose properties varied according to the sensory cue condition (see B. for more details). The dots were visible within a circular aperture in a flat surface positioned at the fixation distance of the display. “Virtual reality headset”: Seated observers viewed the stimuli in an Oculus DK2 head-mounted display. Their head movements were tracked (6 degrees of freedom) and recorded in all conditions. Depending on the experimental condition, the virtual scene updated according to the head movements. “Percept”: Observers fixated the center of the circular aperture. The random dot stimulus would appear at fixation and appear to move either towards or away from the observer for 250 ms before disappearing. Observers indicated the perceived direction of motion by button press. Observers were given unlimited time to respond. Subsequently, both visual and auditory feedback were provided. **B**. Illustration of the four sensory cue conditions tested in the motion task. In the Monocular cues condition, the dot stimulus was randomly presented to one eye on each trial. The dots changed in size and density according to their motion direction. In the Binocular cues condition, binocular disparity and interocular velocity change cues were present, while dot size and density were held constant. In the Combined cues condition, binocular disparity and interocular velocity change cues as well as dot size and density cues were present in the stimulus. Finally, in the Full VR condition, all of the cues in the Combined condition were present, as well as motion parallax cues due to head-motion contingent updating of the display.

The display was calibrated using standard gamma calibration procedures. Minimum and maximum display luminances were 0.01 cd/m_2_ and 64.96 cd/m_2_, respectively. The experiment was controlled by MATLAB and the Psychophysics Toolbox [18]-[20] on a Macintosh computer and projected on the display of the DK2 headset. During the psychophysical task portion of the study (see next section, “3D motion direction discrimination task”), observers used a keyboard to initiate trials and make responses.

### 2.3. Experimental Procedure

Each observer completed a single 1-hour experimental session. After observers gave informed consent, their static stereoacuity was measured using the Randot Stereotest (Stereo Optical, Chicago, IL). All but two observers met the criterion of reaching level 5 (70 arc sec) on its graded circles test. The remaining two observers achieved a level of 4 (100 arc sec), but were included in subsequent data analyses after statistical tests demonstrated that their performance did not differ from the remaining sample. The inter-pupillary distance (IPD) was then measured for each observer using a pupillometer (Essilor Instruments, USA), providing measures in half-millimeter increments. Observers next completed the Simulator Sickness Questionnaire (“baseline SSQ”; [21]). Upon completion of the questionnaire, observers underwent a brief calibration procedure in which the DK2 headset was calibrated for their IPD and height. They were then introduced to the experimental task and completed 50 practice trials (see “Motion task” section below for more details) in the presence of the experimenter.

The experiment then began with the sensitivity assessment, which we describe in more detail below. Observers completed four 2.5-minute blocks of the motion task in a random, counterbalanced order across observers. Observers took brief breaks between these blocks, during which they completed the SSQ (“post task”). After the final block and SSQ, observers entered the motion sickness phase of the experiment. They watched up to 22.5 min of stereoscopic video content with the option to quit if the experience became intolerable. Upon completion of the video content (or quitting the viewing), observers completed another SSQ (“post video”). In the final five minutes, observers were asked to complete a brief questionnaire reporting on their experience with motion sickness and virtual reality prior to our experiment, and they were debriefed about the study. Prior to leaving, they completed a final SSQ (“end of session”). During all phases of the experimental procedure, observers remained seated. No restraints (i.e., forehead or chin rests) were used.

### 2.4. Motion Task

The sensitivity assessment was based on observers’ performance on a 3D motion direction discrimination task. Observers judged the motion direction of a set of 12 white dots 0.2 cm in diameter at the 1.2 m fixation distance (**Fig. 1A**, “Virtual display” panel). The dots appeared at the center of a visual scene and moved toward or away from the observer at ∼96 cm/s for 0.25 s before disappearing. These world-based stimulus parameters translate to a dot diameter of 0.1 deg (or 0.9 pixels) at the fixation distance. Dots moved at 1.2 deg/s. When a dot reached a disparity of ±0.15 we flipped its disparity sign and assigned a new *x* and *y* position. In this way, the stimulus appeared as a volume of dots centered on the fixation plane where each dot moved continuously toward or away from the observer. Note that in all but the Binocular Cues condition (see below), dot size and density changed in a manner consistent with projective geometry. As illustrated in the Supplementary Materials videos, changes in dot size were probably not very noticeable to observers given the resolution of the display, but changes in dot density could be clearly seen.

Motion coherence was manipulated by varying the proportion of signal to noise dots. For each trial, we pseudo-randomly selected a motion coherence level from [0% 16.67% 50% 100%] coherence for 13 of the observers, and from [16.67% 50% 100%] for the remaining 82 observers.

On each stimulus frame, we randomly selected a subset of dots as signal dots, which moved coherently, either toward or away from the observer (perpendicular to the screen). The remaining dots (noise dots) were given random *x, y*, and *z* coordinates. Signal and noise dots were selected on a frame-by-frame basis to help prevent observers from tracking the direction of motion of individual dots. Direction of motion (i.e., “toward” or “away”; see **Fig. 1A**, “Percept” panel) was chosen pseudo-randomly on each trial.

Multiple visual cues signal motion in depth [22]-[24]. We aimed to quantify observer sensitivity to each cue by manipulating the available cues in the dot motion stimulus. We tested sensitivity in four conditions: *Monocular cues* (dot size and density changes were presented, but binocular cues were eliminated by presenting the stimulus to one eye only), *Binocular cues* (containing binocular disparity and inter-ocular velocity differences, but monocular cues were eliminated by keeping dot size and density constant), *Combined cues* (containing both the monocular and binocular cues), and *Full VR* (containing the combined cues as well as motion parallax cues due to head movement) (see **Fig. 1B** and see **Supplemental Material** for videos illustrating the four cue conditions). It is important to note that in the Monocular cues condition, the dots were presented to one pseudo-randomly chosen eye on each trial.

The motion stimuli were presented in the center of a virtual room (3 m in height, 3.52 m in width, and 3.6 m in depth). While this room served no function during the actual experiment, it helped observer immersion during initial instruction. The virtual walls, ceiling, and floor were all mapped with different tiled textures to facilitate better judgment of distances throughout the virtual space and judgment of the stimulus motion trajectories. The room contained a surface (i.e. wall) that was positioned at the display’s focal distance (1.2m from the observer). The wall was textured with a 1/f noise pattern that aided accommodation and vergence. Stimuli were presented within a 3 deg radius circular aperture located in the center of the wall with the dots confined to the central 2.4 deg to prevent occlusion by the aperture’s edge. Thus, dots appeared to move through an aperture in a wall, and either approached or receded from the observer depending upon the pseudo-randomly selected motion for that trial. A small (0.04 deg) white fixation point was visible in the center of the aperture at all times except when a dot motion stimulus was presented. All stimulus elements were anti-aliased to achieve subpixel resolution.

Observers were instructed to report the dot motion direction. Observers indicated the direction of dot motion by pressing the up arrow key on the keyboard for receding motion and the down arrow key for approaching motion. In recent work, feedback was shown to be critical for the recruitment of sensory cues in VR displays, especially binocular and motion parallax cues to motion-in-depth [25]. Likewise, to encourage recruitment of the sensory cues in the different conditions in the current study, observers received auditory feedback (a “cowbell” sound on correct trials and a “swish” sound on incorrect trials) as well as visual feedback (behavioral performance up to and including the current trial in terms of percent correct was displayed at the fixation point). If the most recent response was correct, the performance was displayed in green; if incorrect, in red.

During stimulus presentation, observers were asked to keep their head still and maintain fixation. In all but the Full VR condition, head movement had no effect on the display so that it appeared to the observer that the virtual environment was “glued” to the head. In the Full VR condition, the visual display updated according to head movement, so that it appeared that the observer was present in a stationary immersive virtual environment.

Observers completed the task in four 2.5-minute blocks after completing 50 practice trials with feedback in the Full VR cue condition. Observers that were presented with four coherence levels completed 84 trials, and observers that were presented with three coherence levels completed 90 trials with each of the blocks. All observers completed four blocks in a randomized, counterbalanced order. Each block contained stimuli from one of the four cue conditions (Monocular, Binocular, Combined, or Full VR). Between blocks, observers took short breaks during which they removed the VR headset and completed the Simulator Sickness Questionnaire (see “Quantifying motion sickness” section below).

### 2.5. Video Content

Observers viewed up to four stereoscopic videos [17], totaling ∼22.5 min in the VR headset, played in Windows Media Player. The four videos increased in level of intensity: (1) a 5:34 min, first-person video of a car driving through mild traffic, (2) a 3 min first-person computer-generated (CG) video of a fighter jet flying through a canyon, (3) a 6:26 min first-person video of a drone flying through a parking lot, and (4) a 7:19 min first-person video of a drone flying around a bridge (see **Fig. 2** for stills from the four videos). All observers watched the videos in the same order. Observers were told they could stop viewing the videos if and when the experience became intolerable. The total viewing time was recorded for each observer.

**Figure. 2.**
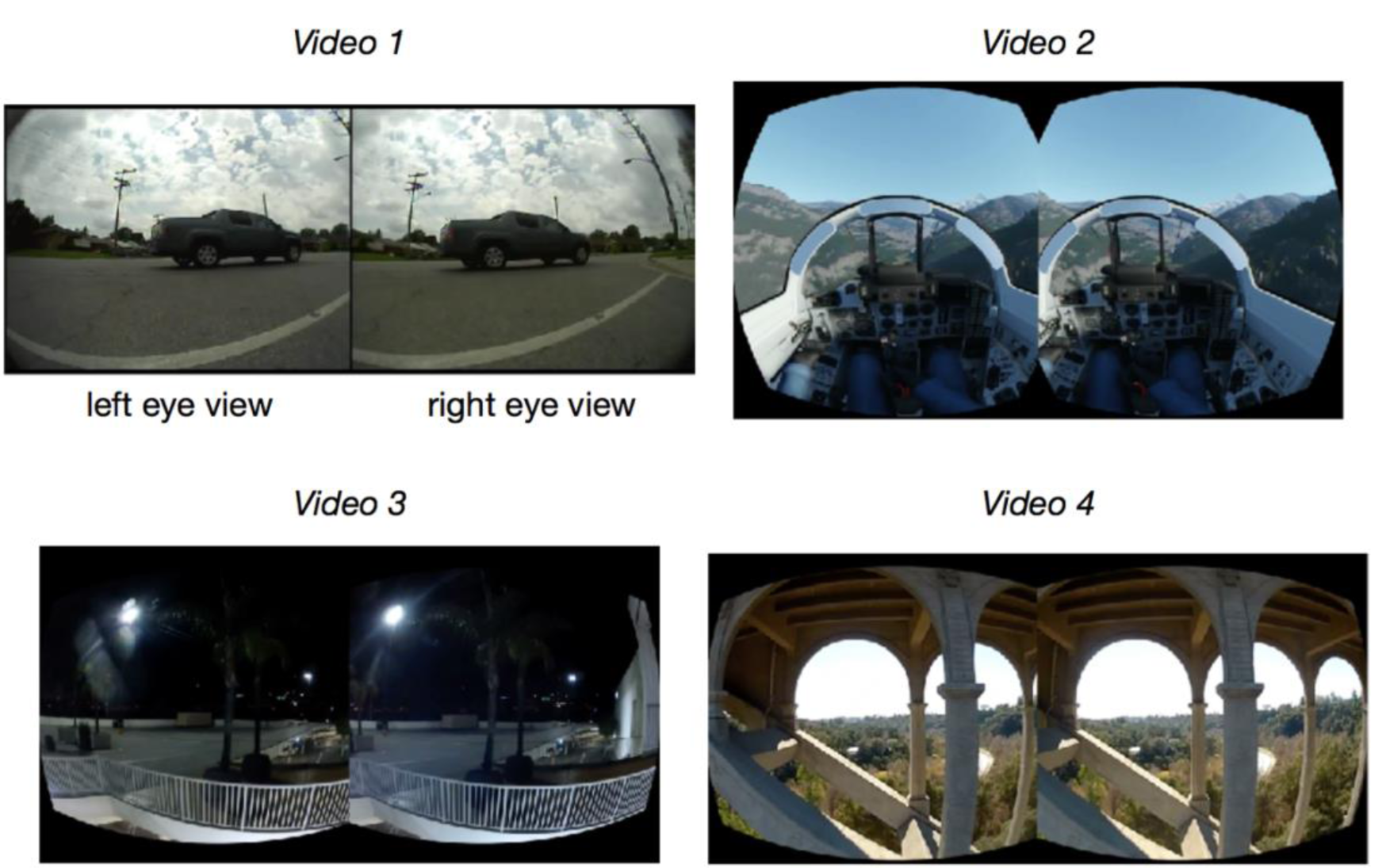
Stills from motion sickness inducing videos. After completing the four blocks of the motion task, all observers viewed up to four videos while wearing the Oculus DK2 head-mounted display in the same order (up to 22.5 minutes). The videos increased in intensity, and observers were given the option to quit if viewing became intolerable. All observers, whether they stopped the video viewing early or not, completed the Simulator Sickness Questionnaire (SSQ) to indicate the severity of motion sickness symptoms.

### 2.6. Data Analysis

#### 2.6.1. Quantifying Sensitivity

For each cue condition, we calculated the proportion of ‘toward’ responses as a function of direction and motion coherence. Standard error of the mean (SEM) was calculated based on the binomial distribution of the (toward/away) responses. We fit the proportion of toward responses *g(x)* as a function of direction and motion coherence (*x*) with a cumulative Gaussian using nonlinear least squares regression using the lsqcurvefit function in MATLAB:

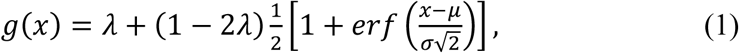

where *x* is the directionally-signed motion coherence, μ is the estimate of observer bias, σ reflects the precision of the responses for the respective sensory cue condition, and λ is the lapse rate. To stabilize fits when precision was low, we enforced a bound of ±.5 on μ and bounds of [0.01 100] on σ. The fitting procedure assumed a lapse rate of 2%. Sensitivity was computed as 1/σ.

We computed median fit parameters using a bootstrap procedure to ensure stable estimates. We resampled the “toward”/“away” response data with replacement 1,000 times for each observer. We then fit a psychometric function to each resampled data set. Finally, we obtained the median parameter estimates from the fits.

To determine if the performance of each observer for each cue condition was different from chance, we simulated performance of an observer who responded randomly on each trial for 90 total trials for three coherence levels and 84 trials for four coherence levels. We fit psychometric functions to 10,000 simulated data sets and obtained the sensitivity estimate from each. At the 95% confidence level, the upper sensitivity bound was .49/.43 for the conditions with four and three coherence levels, respectively. If an observer’s performance did not exceed these bounds, (i.e., perform above chance level) in any of the four conditions, the observer was excluded from further analyses (n = 8).

#### 2.6.2. Quantifying Motion Sickness

To quantify motion sickness, observers completed the SSQ at six separate time points during the experimental session (see “Experimental Procedure” above). Observers rated the severity of 16 symptoms as “none”, “slight”, “moderate”, or “severe”, which were then numerically scored as 0, 1, 2, and 3, respectively. The symptoms form three subscales: (1) nausea (N) ranging from 0 - 200.34, (2) oculomotor disturbances (OD) ranging from 0 - 159.18, and (3) disorientation (D) ranging from 0 - 292.32. The severity of symptoms on each of the three scales was computed via standardized formulas (see [21]), which were then combined using a final formula to produce an overall (“Total”) sickness score ranging from 0 - 235.62. Larger scores correspond to more severe symptoms on all scales. Although the sickness scores were computed for each of the six questionnaires completed by each observer during the experimental session, we were primarily interested in the effects of the video viewing. To quantify the impact of video viewing on sickness levels, we computed the change in motion sickness from before the video viewing (based on the “post task” SSQ) to after the video viewing (“post video” SSQ).

#### 2.6.3. Quantifying Head Movement

We used the head-tracking capabilities of the virtual reality device to measure head movement during the assessment of visual sensitivity. Head movements during the task were relatively small due to the presentation of the stimulus at fixation for a brief time - we therefore refer to these small head movements as “head jitter” [25]. We analyzed translational head jitter and rotational head jitter based on the 6 DoF head tracking built into the DK2 headset. For each block of the motion task, a single continuous head trace was saved, containing the 4×4 model view matrix for each eye at every screen refresh (75 Hz or ∼13.33 ms). We inverted the model view matrix and determined the “cyclopean” view matrix at each time point based on the midpoint between the two eyes’ views. From these traces, we extracted the time points that corresponded to each individual trial in order to analyze the head movement on a trial-by-trial basis. No additional transformations were applied.

To quantify translation, we computed the head’s path length through 3D space (‘translational jitter’) for each trial. We path-integrated the translation of the head by summing the Euclidean distance between each consecutive head position obtained from the X, Y, and Z components of the “cyclopean” view matrix. Point-to-point estimates ≥ 0.002 m (which corresponded to a velocity ≥ 0.15 m/s) were excluded because they were unrealistically large and likely reflected tracking errors (∼9.5% of all time points across all observers, conditions, and trials). Thus, when an erroneous tracking time point was identified, the path integration ignored that point. Because the distributions of translational jitter were typically positively skewed, we computed the median translation for each observer. Average translational jitter was then computed across observers.

Similar methods were used to quantify rotation. We first computed the total angular distance that the head rotated in 3D space on each trial (‘rotational jitter’). To do so, we extracted the rotation components (i.e., the first three rows and columns) from the 4×4 “cyclopean” view matrix *M. M* was then decomposed to determine the amount of rotation about each axis in the following order: y (yaw), z (roll), and x (pitch). To calculate the total rotation, the observers’ orientation at the start of the trial was represented by the vector (0, 0, 1), which corresponded to the observer looking straight ahead. Following time point 1, the direction vector at each time point was calculated by computing the dot product of *M* and the starting vector (0, 0, 1). Total rotational jitter was computed by summing the total head rotation between every two adjacent time points (i.e., the absolute angle between two successive vectors). Point-to-point estimates of rotation that were ≥ ∼28.5 arcmin (which corresponded to an angular velocity of ∼36 deg/s) were excluded (<1% of all time points across all observers, conditions, and trials). As with the computation of translational jitter, when an erroneous tracking time point was identified, the path integration ignored that point. Rotational jitter distributions were typically positively skewed, so we computed the median rotation for each observer. Average rotational jitter was then computed across observers.

### 2.7. Statistical Analysis

To assess changes in motion sickness SSQ scores were obtained at three different timepoints (baseline, post-task and post-video). We subsequently computed the difference between baseline and post-task scores, as well as post-task and post-video scores for each observer and computed the significance of these differences using the Wilcoxon signed rank test. Since we evaluated the Total SSQ scores as well as the scores on each of the three SSQ subscales these tests were carried out at the Bonferroni-corrected alpha level of .0125.

The relationship between sensitivity in each stimulus condition and motion sickness due to video viewing were quantified through an analysis of variance (ANOVA) evaluated on general linear model fits to the individual subject data for each of the sensory cue conditions with sensitivities (1/σ_*cue*_) included as a fixed effect and subject included as a random effect, specified as ΔSSQ ∼ 1/σ_*cue*_ + (1 | Subject). Individual sensitivity values that were three standard deviations beyond the mean in each of the cue conditions were excluded from the analysis, yielding: n = 95, 93, 94, 94 data points included in the model for the Full VR, Combined, Monocular, and Binocular, respectively. A Bonferroni-corrected alpha level of .0125 was used to test for significance of the four relationships. Effect size is reported as **Ω**^2^.

The role of sex in the relationship between sensitivity to the cues in the Full VR condition and motion sickness due to video viewing was evaluated through an ANOVA evaluated on general linear model fits to the individual subject data with sensitivity to the Full VR condition (1/ σ_*FullVR*_) and sex included as fixed effects along with their interaction and subject included as a random effect. Significance of the main effects and the interaction was evaluated at the alpha = .05 level.

Patterns of head jitter were analyzed over time. For each observer, head jitter was averaged for each trial over the four blocks of the motion task (i.e., the sensitivity assessment portion of the experiment), giving a within-subject mean head translation (in mm) and within-subject mean head rotation (in arcmin) for each trial. We fitted linear, quadratic, and power models to the between-subject mean head translation and between-subject mean head rotation across trials, with the first 5 trials omitted to ensure stable behavior at the start of the trial. An AIC model comparison indicated that the quadratic model best-characterized the pattern of head translation and rotation across trials and subjects. We then carried out two multiple quadratic regressions, one for translational head jitter and one for rotational head jitter. These models tested for an effect of average observer sensitivity to the sensory cue conditions on head jitter, controlling for trial (i.e., time spent in the device) with subject included as a random effect. N = 8075 total data points per head jitter type were supplied to the model, however, outliers that were 3 standard deviations beyond the mean at each time point (i.e., trial) were excluded for consistency with other analyses (∼1% & ∼2% of all data points for translational and rotational head jitter, respectively). This exclusion did not change the overall results or their interpretation. Significance of the effect of sensitivity was evaluated at the alpha = .05 level.

## 3. Results

### 3.1. Variability in Sensitivity to 3D Motion Cues in VR

We first assessed sensitivity to 3D motion cues in virtual reality. Each observer judged the direction (toward/away) of a cloud of dots moving with variable coherence levels. We fit a cumulative Gaussian to the observer’s performance and used the inverse of the standard deviation (1/σ) as an estimate of the observer’s sensitivity. Each observer’s motion sensitivity was tested in four cue conditions (Monocular, Binocular, Combined, and Full VR). Combined stimuli contained both monocular and binocular cues, and Full VR stimuli contained the Combined condition cues as well as motion parallax cues.

We found considerable variability in sensitivity to the different sensory cues (**Fig. 3**). Cue sensitivity varied both within and across observers. On average sensitivity was greatest when more cues were available (Full VR and Combined Conditions), and smallest when fewer cues were available (Monocular and Binocular Conditions), and binocular cue sensitivity was generally weakest. However, observers with lower sensitivity in one sensory cue condition were not necessarily those with lower sensitivity in the other conditions. Importantly, variability among observers *within* each sensory cue condition was larger than the variability in sensitivity *between* the four cue conditions. These effects do not appear to be related to stereoacuity as Randot performance did not predict sensitivity in any of the cue conditions (p > .05 for all conditions).

**Figure. 3.**
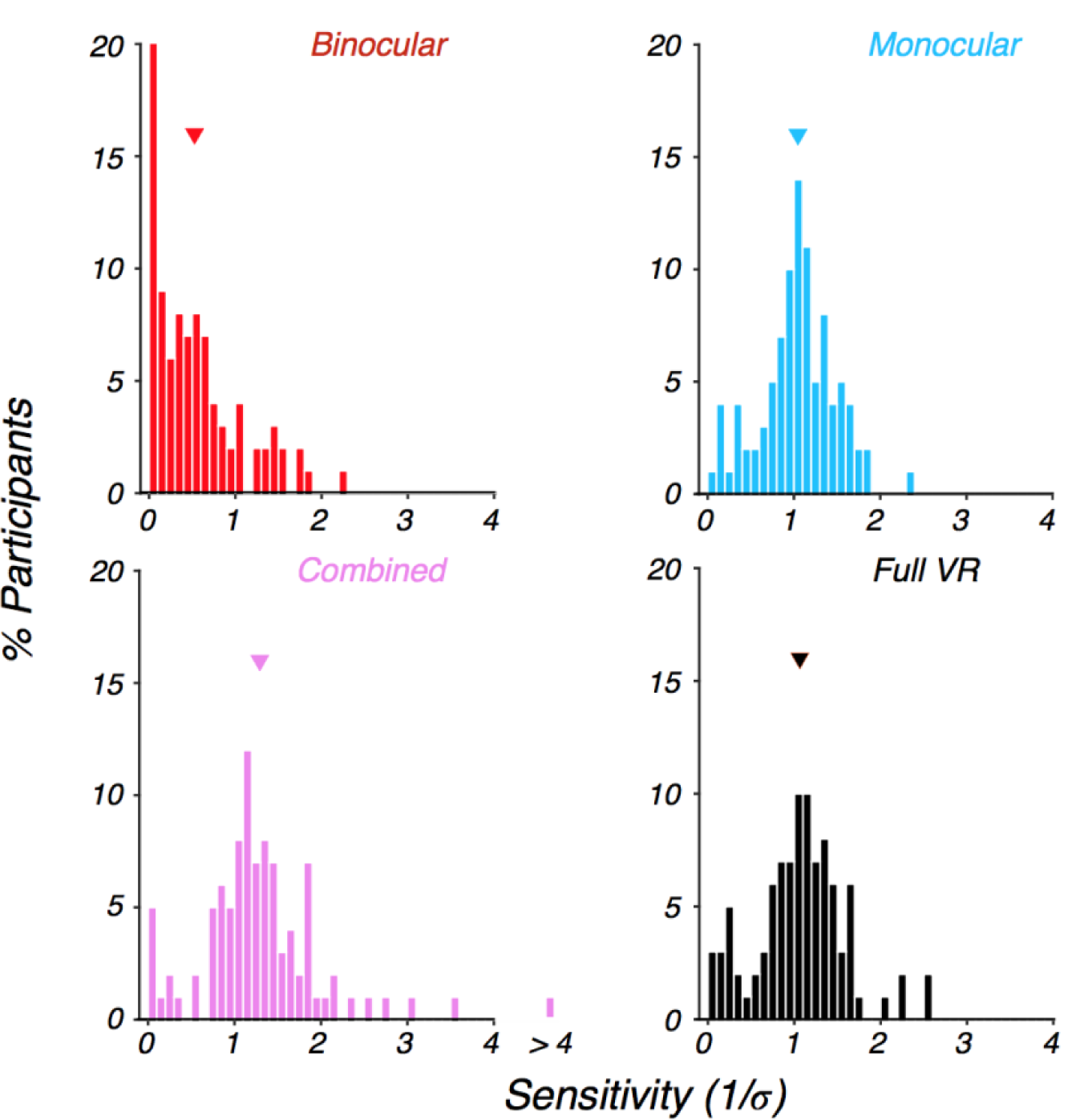
Sensitivity to 3D motion cues varies across observers. On average sensitivity is greatest when more cues are available (Full VR and Combined Conditions), and smallest when fewer cues are available (Monocular and Binocular Conditions), with binocular cue sensitivity being particularly poor. However, variability among observers within each sensory cue condition was considerably greater than the variability in sensitivity between the four cue conditions, indicating considerable inter-observer sensitivity differences to each cue. Each panel reflects the distribution of sensitivity to the particular cue condition across n = 95 observers. Bars in the histograms correspond to the % of participants falling in each sensitivity bin, and the triangles correspond to the between-subject mean sensitivity for the condition.

### 3.2. VR Video Content Induces Motion Sickness

We next assessed the susceptibility to motion sickness in the same observers using the Simulator Sickness Questionnaire (SSQ; [21]). This questionnaire was developed to quantify the symptoms most commonly experienced by users of virtual reality systems and has been normed to provide scores on three categorical scales. Larger scores indicate more intense motion sickness symptoms. Observers completed the SSQ at several points in time throughout the study (see Methods for more details): 1. after consenting to participate in the study, prior to any VR exposure (“baseline”); 2. immediately after the motion task, prior to viewing the video content (“post task”); 3. immediately after viewing the video content (“post video”).

Observers generally reported little sickness at the beginning of the study (**Fig. 4**, blue bars). Increases in motion sickness symptoms were reported after completion of the motion task (red bars). Wilcoxon signed rank tests of the pre- and post-task SSQ ratings indicated that ratings were significantly higher for the post-task Total SSQ and the three SSQ subscales (p < .001 for all tests). Larger increases in motion sickness were observed post-video viewing, producing moderate levels of motion sickness on average. Wilcoxon signed rank tests of the pre- and post-video SSQ ratings indicated that ratings were significantly higher for the post-video Total SSQ and the three SSQ subscales (p < .001 for all tests), confirming that our manipulation of motion sickness had its intended effect (yellow bars). Of note, as with the results of the sensitivity assessment (i.e., performance in the motion task), there was considerable variability across observers in the intensity of motion sickness symptoms throughout the study, with some individuals appearing highly sensitive to the manipulation and others apparently insensitive to it.

**Figure. 4.**
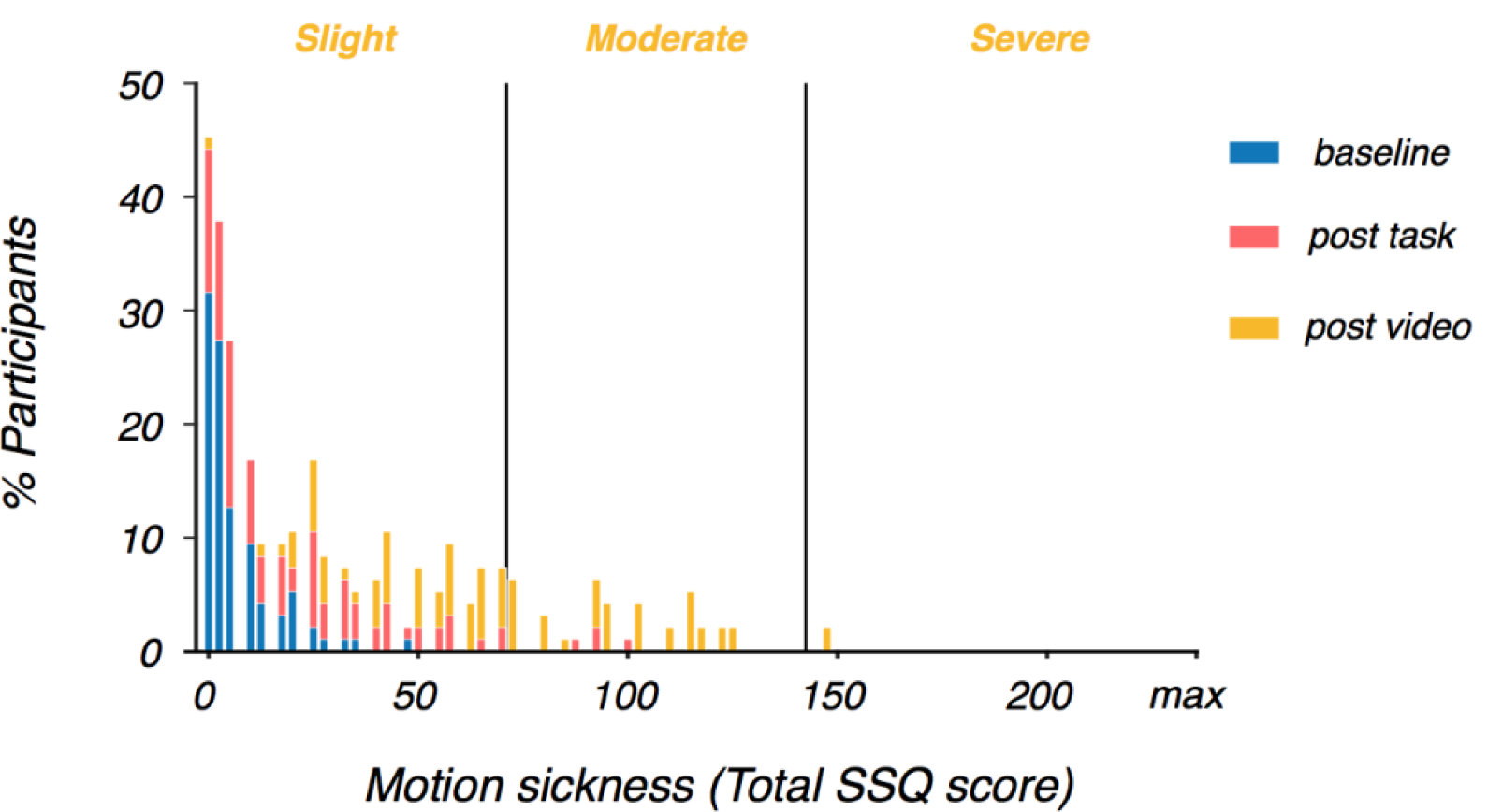
VR video viewing increased motion sickness. Prior to any VR exposure in the laboratory (baseline - blue bars), observers reported minimal sickness symptoms. Post motion task (i.e., the cue sensitivity assessment - red bars), observers reported slightly elevated sickness symptoms on average. Post video viewing (orange bars), observers reported moderate sickness symptoms on average, as expected. In the analyses reported below, we focused on the change in sickness symptoms directly pre and post video viewing (i.e., post video - post task). The maximum attainable score on the Total SSQ scale is 235.62. See Methods for details.

The increased levels of motion sickness with video viewing were not unexpected given the sensory cue conflicts in the video content. In particular, although care was taken to ensure that the HMD was tailored to the inter-pupillary distance (IPD) of each observer, the binocular disparity in the video content was fixed according to the disparity of the original recording. In addition, the video content provided motion parallax information consistent with the recording camera’s movement, not the observer’s head movement.

### 3.3. Sensitivity to 3D Motion Cues Predicts Motion Sickness

We predicted that observers with greater sensitivity to sensory cues would experience more severe motion sickness. Specifically, we hypothesized that sensory cue conflicts give rise to motion sickness, and observers with high sensory sensitivity would be more likely to detect cue conflicts while viewing the VR videos. Thus, when considering the relationship between the motion sickness related to video viewing and sensitivity to the sensory cues, we expected a positive relationship. We found the expected positive relationship in the Full VR condition (*F*(1,93) = 14.21, *p* < .001, **Ω**_2_ = .1302; see **Fig. 5)**. We did not find a significant relationship between cue sensitivity and motion sickness in any of the other conditions (*p* > .0125, the Bonferroni-corrected alpha-value; see **Table 1**).

**Table 1:** Nausea and disorientation in VR are predicted by sensitivity to motion cues in the Full VR condition. Entries correspond to the p-values of the relationships between sensory cue sensitivity in each cue condition and motion sickness due to VR video viewing. Bold p-values with an asterisk denote significance at the Bonferroni-corrected alpha level = .0125. The total sickness score is derived from a combination of the scores on the three separate sub-scales: Nausea, Disorientation, and Oculomotor discomfort. The significant relationship between the total sickness score and sensitivity to the cues in the Full VR condition are primarily driven by Nausea and Disorientation scale symptoms. The trend towards a relationship between the total sickness score and sensitivity to the cues in the Monocular condition may be primarily driven by Oculomotor discomfort arising from the dot stimulus being visible in only one eye on each trial.

| Cue condition | Motion sickness (SSQ) |  |  |  |
| --- | --- | --- | --- | --- |
|  | Total sickness | Nausea | Disorientation | Oculomotor Discomfort |
| Monocular | $p = .044$ | $p = .073$ | $p = .279$ | $p = .029$ |
| Binocular | $p = .879$ | $p = .314$ | $p = .636$ | $p = .906$ |
| Combined<br>(Monocular + Binocular) | $p = .125$ | $p = .039$ | $p = .556$ | $p = .227$ |
| Full VR<br>(Combined +<br>Motion parallax) | **$*p = .0003$** | **$*p = .00002$** | **$*p = .003$** | $p = .041$ |

*\*denotes significance at the Bonferroni-corrected alpha level of .0125*

**Figure. 5.**
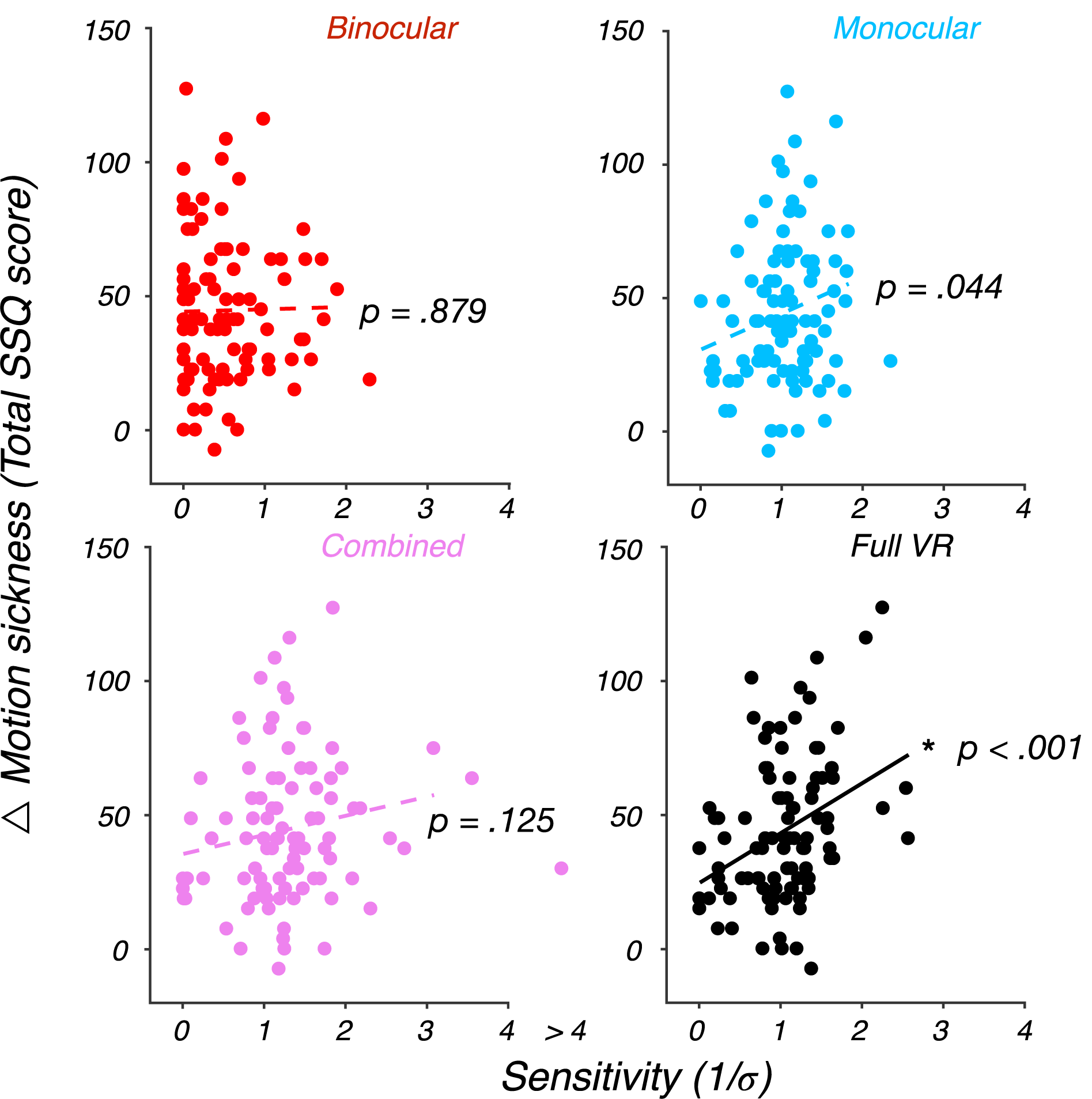
Sensitivity to motion cues in the Full VR condition predicts motion sickness. For each of the four sensory cue conditions, we computed the relationship between sensitivity to the sensory cues and severity of motion sickness due to video viewing. Solid line denotes a significant relationship at the Bonferroni-corrected alpha level = .0125. The relationship is significant only for the Full VR condition. Given that the Full VR condition is the only of the four sensory cue conditions that contains motion parallax cues, this result suggests that sensitivity to motion parallax information in particular predicts sickness due to video viewing where motion parallax cues are unavailable.

This relationship was specific to two of the three SSQ sub-scales. Sensitivity to the sensory cues in the Full VR condition was highly-correlated with Nausea scale symptoms (*F*(1,93) = 19.79, *p* < .001, **Ω**_2_ = .1724) and Disorientation scale symptoms (*F*(1,93) = 9.21, *p* = .003, **Ω**_2_ = .0884). No relationship was identified between Full VR sensory cue sensitivity and Oculomotor Discomfort scale symptoms (*p* > .0125, the Bonferroni-corrected alpha-level). We did not find significant relationships between sensitivity to the sensory cues in the Monocular, Binocular, and Combined conditions and these sub-scales (*p* > .0125 in all cases, see **Table 1**).

Thus, we found a strong relationship between motion sickness severity and sensory sensitivity in the Full VR condition, which contained motion parallax cues. By contrast, we find little evidence for such a relationship in any of the other conditions, which did contain various other cues to stimulus motion, but did not contain motion parallax cues. Taken together, these results confirm the role of cue conflicts in motion sickness, suggesting that observers who are more sensitive to visual motion parallax cues are more likely to develop sickness symptoms in VR.

### 3.4. No Relationship Between Sex and Motion Sickness

The above analysis indicates that sensitivity to sensory cues, particularly to motion parallax cues, plays a critical role in motion sickness. Previous work has revealed sex differences in susceptibility to motion sickness as well. Specifically, women are thought to be more susceptible to severe motion sickness due to greater postural instability and larger postural sway in non-VR [26], as well as VR environments [27].

We tested for a relationship between sex and motion sickness in addition to the sensitivity to the cues in the Full VR condition. However, we did not find a significant role of sex in our data (*F*(1,91) = 0.83, p = .36), nor an interaction between sex and sensitivity in motion sickness (*F*(1,91) = 2.61, *p* = .11). Finally, the relationship between sensitivity and sickness reported above remained significant when accounting for the sex of the observer in our model (*F*(1,91) = 14.91, *p* < .001; **Fig. 6**).

**Figure. 6.**
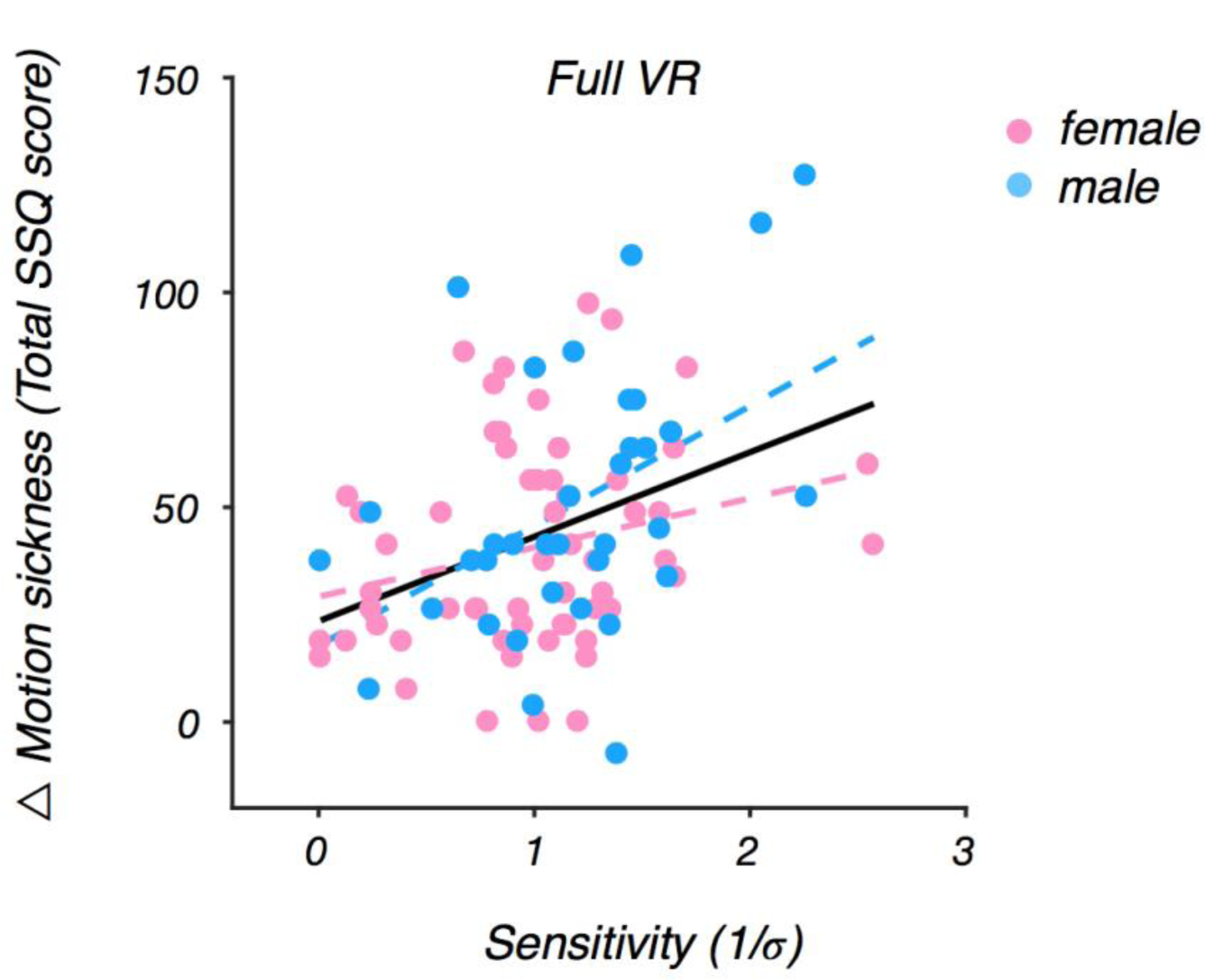
Motion sickness is predicted by visual sensitivity in Full VR but not sex. The plot depicts the same data as in Fig. 5 - Full VR with sex of the observer denoted. Solid line denotes significance at the alpha = .05 level. The relationship between sensitivity and sickness with effect of sex removed remains significant (solid line), and the effect of sex is not significant. The dashed lines correspond to the sensitivity - sickness relationship for female (pink) and male (blue) observers.

A possible source of discrepancy between current results showing no effect of sex and previous reports may relate to inter-pupillary distance (IPD; [16]). Previous studies have largely relied upon a default IPD when presenting experimental stimuli. Default IPDs of stereoscopic stimuli are typically set to 63-64 mm. In the current study, however, we tailored the device to the IPD measurements taken for each observer at the start of the experiment.

Why might this be a source of the difference in sex effects? Consideration of the distribution of IPDs in our sample (see **Fig. 7**) reveals that the average IPD for males is closely matched to the default device IPD of 64 mm. We should note, however, that the default IPD still misses the mark for many of the males in our sample. The situation is worse for females, for whom the average IPD is nearly 5 mm smaller than the default IPD. Mismatches between device and observer IPD will inevitably introduce cue conflicts, which will lead to motion sickness. Our results suggest that tailoring the IPD of the display to the individual may reduce motion sickness - that is, ensuring that the IPD of the device is matched to the IPD of the observer will reduce some sources of cue conflicts and will likely eliminate the sex differences reported in previous work (see also [16]).

**Figure. 7.**
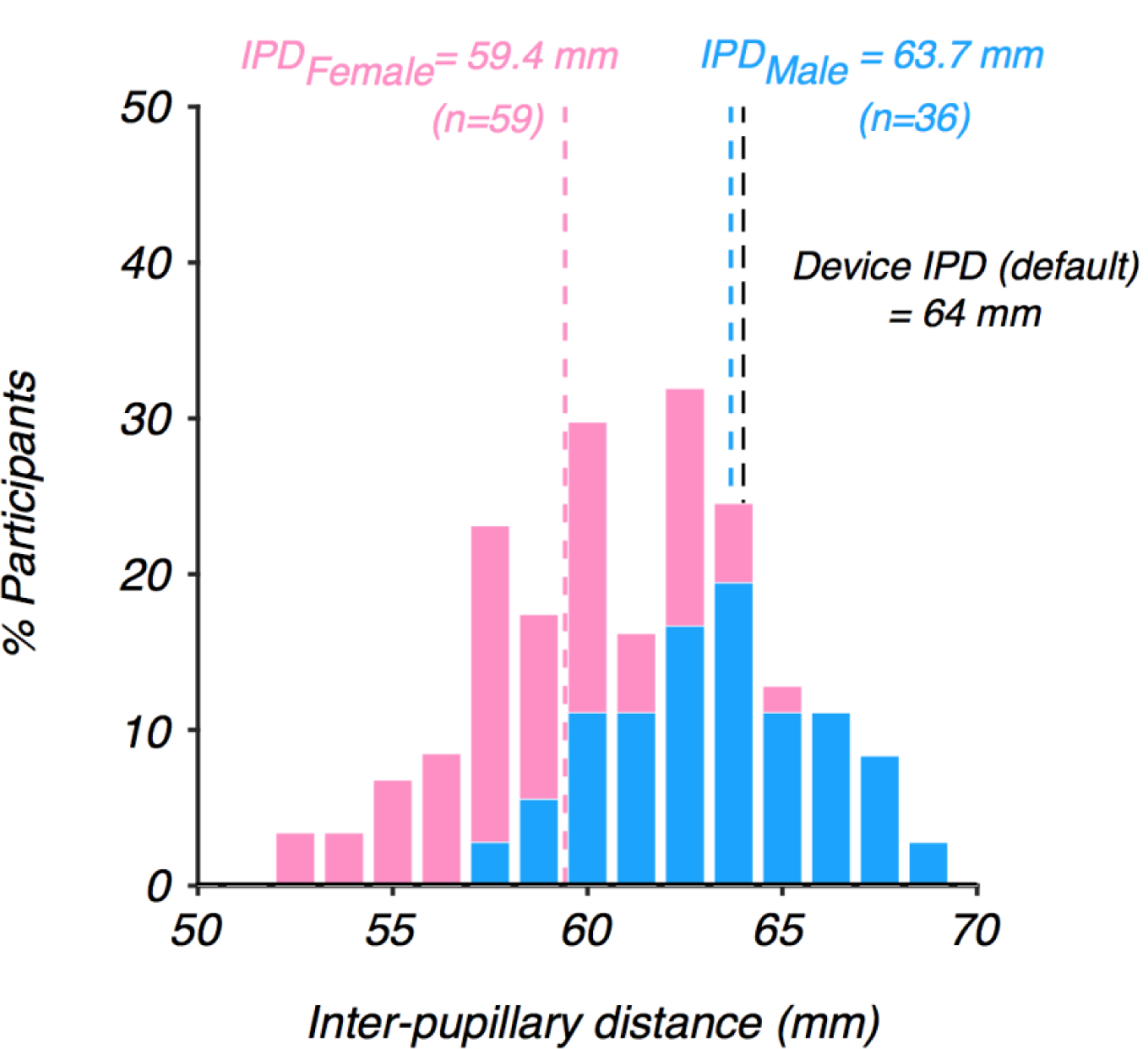
Inter-pupillary distance (IPD) for the sample of females (pink bars) and males (blue bars) in our experiments. The average male IPD is well-matched to the default IPD of the Oculus DK2, whereas the average female IPD is approximately 5 mm smaller than the default.

One might assume that larger IPDs imply greater sensitivity to binocular cues and hence, that IPD *per se* is an important factor in motion sickness. This assumption is not backed up by our data - no relationship was found between IPD and average sensitivity *(F*(1,93) = 0.49, *p =* .49). Moreover, although there was a trend towards individuals with larger IPDs reporting more severe levels of motion sickness due to video viewing, this relationship also did not reach significance *(F*(1,93) = 3.82, *p* = .054). Finally, no relationship was found between observer height and motion sickness *(F*(1,93) = 0.64, *p* = .43). Therefore, the large variability in sensitivity to the 3D motion cues measured in our sample must lie in subsequent processing of motion-in-depth signals, not physical characteristics such as IPD or height *per se*.

### 3.5. Observers Reduce Head Movements to Regulate Motion Sickness

If motion sickness is caused by cue conflicts, a useful observer strategy would be to actively eliminate cue conflicts when motion sickness occurs. Since motion parallax-based conflicts appear to be the primary source of motion sickness in VR, observers could eliminate conflict by reducing head movement. This line of reasoning predicts that as individuals start to experience discomfort, they may suppress their head movement. To test whether this strategy is adopted by observers, we analyzed the head movement data collected during the four blocks of the motion task. Note that in three of those blocks, motion parallax cues were absent from the display and were thus in conflict with the parallax cues the observer should expect when they moved their head.

Because the stimuli were presented at fixation for a brief duration (250 ms), observers’ head movements were small (on the order of millimeters and arcmins), and we refer to them as “head jitter”. We first analyzed head jitter over the course of an experimental block to determine whether there is evidence of head jitter suppression. We found that on average across observers and experimental blocks, head jitter exhibited a U-shaped pattern: both the magnitude of translational and rotational head jitter declined before rebounding later in the experimental block (see **Fig. 8**). We interpret this pattern as the predicted suppression of head jitter to mitigate motion sickness symptoms, eventually transitioning to a “release” in head jitter once the end of the experimental block is in sight.

**Figure. 8.**
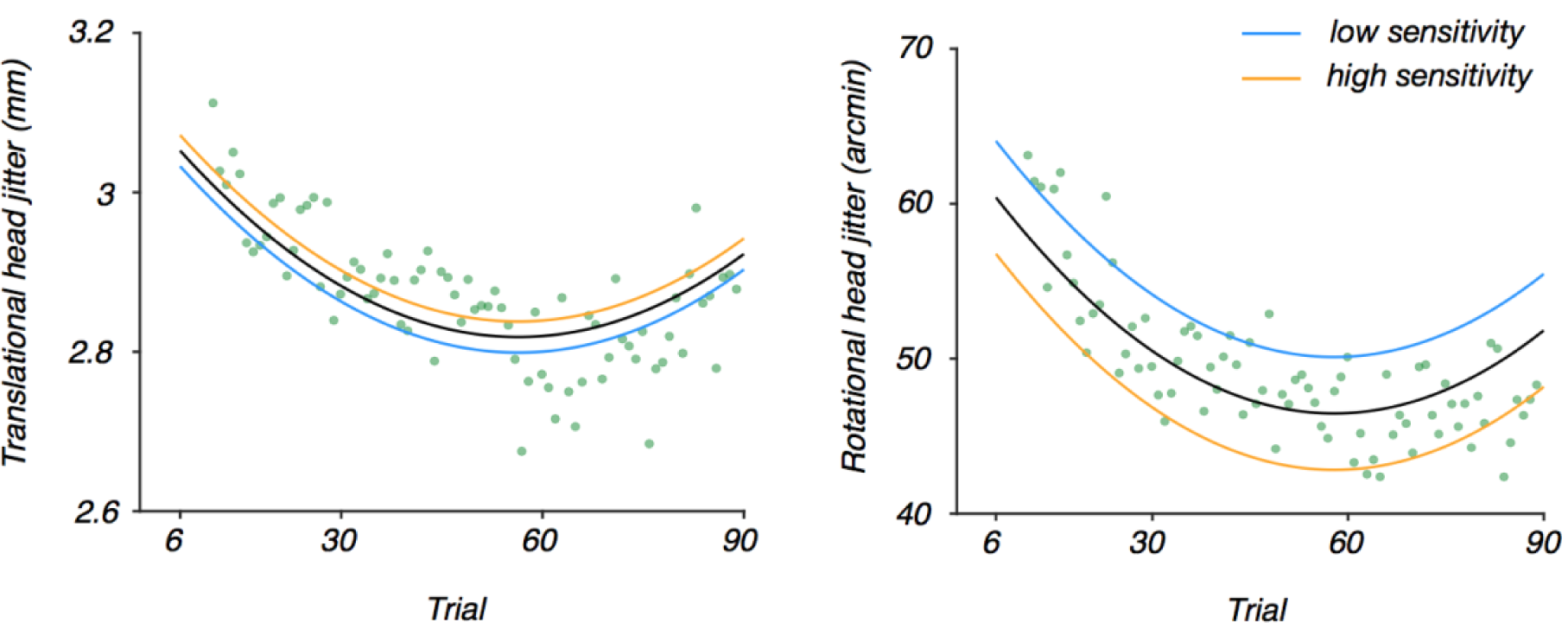
Modulation of head jitter across trials. For each observer, the average 3D translational head jitter and average 3D rotational head jitter were computed over the four blocks across trials. Data points depict the between-subject average of 3D translational head jitter (left plot) and average 3D rotational head jitter (right plot) across trials. Solid black lines correspond to the quadratic fit to the individual subject data points across trials. Orange lines correspond to fits for a high sensitivity observer whose sensitivity is one standard deviation above the mean, and blue lines correspond to fits for a low sensitivity observer whose sensitivity is 1 standard deviation below the mean. The quadratic pattern of both head jitter types indicates that observers suppress head movement over time, which may be used as a strategy to mitigate motion sickness symptoms, and then release head movement during later trials when they likely have experienced a reduction in motion sickness symptoms. Sensitivity to 3D sensory cues significantly modulates the degree of suppression of rotational head jitter only, suggesting that head movements along the rotational axes (i.e., yaw, pitch, and roll) contribute more strongly to motion sickness in VR environments.

We next asked whether the degree of suppression was modulated by an observer’s average sensitivity across the four conditions. Specifically, we predicted that observers with greater sensitivity to 3D motion cues would more strongly suppress head jitter. Although translational head jitter exhibited a U-shaped pattern, the effect of sensitivity did not reach significance (*p* = .78). However, we did find a significant effect of sensitivity on rotational head jitter (*F*(1,7779) = 8.671, *p* = .003). For every unit increase in an observer’s sensitivity, rotational head jitter declined by 9.34 arcmin. This, coupled with the fact that head jitter tended to rebound after the initial suppression, suggests that observers dynamically self-regulated their discomfort by reducing their head movement.

## 4. Discussion

An attractive aspect of virtual reality (VR) is that it can be used to present visual stimuli under more realistic viewing conditions. However, VR introduces discomfort for an estimated 25-40% of individuals including motion sickness (e.g., [28]). In the current study, we have provided evidence that such discomfort arises from sensory cue conflicts, in particular, conflicts related to motion parallax cues.

Importantly, a cue cannot be a source of conflict if an observer is not sensitive to that cue. Sensitivity to sensory cues in VR was highly-variable across the large sample of observers we studied. Critically, a robust relationship emerged, whereby the greater an observer’s sensitivity to motion parallax cues, the more severe the motion sickness symptoms. Although motion parallax cues were always present in concert with other cues to 3D motion, the fact that a relationship between cue sensitivity and motion sickness was not evident in any of the other stimulus conditions supports the notion that motion parallax cues in particular drive the conflict between visual and vestibular signals. This finding extends recent work showing that sensitivity to 3D motion cues more generally predicts motion sickness susceptibility [17] and provides targets for future research on causes and mitigation of motion sickness.

Our results also revealed an apparent tendency for observers to self-regulate motion sickness through head movement suppression. Indeed, head movement has previously been implicated in motion sickness. Observers decrease head movement when environments contain conflicting signals, such as in the slow rotation room (SRR; e.g., [4]) or virtual reality [29]. Furthermore, motion sickness is reduced when the observer’s torso or head is restrained [30] although postural precursors of motion sickness still exist under such conditions [31]-[32]. Future work tracking motion sickness over time at more frequent intervals can confirm head movement reduction as a strategy for self-regulation of motion sickness.

Previous work has shown that rotational movements may play a particular role in motion sickness symptoms due to their role in increasing vection, which causes mismatches between visual and vestibular signals in virtual environments (e.g., [33]). Here, we showed that observers in general reduced their head movement in the early portion of each experimental block, followed by a rebound later in the block. Although this pattern was evident in both translational and rotational head movement, rotational head movement suppression was modulated by one’s sensitivity to sensory cues. Thus, we have shown that sickness does not arise from head movement *per se*, but rather is related to an observer’s sensitivity to sensory cues more generally.

Prior work has described the cues that may be in conflict in VR [34] and explored potential links between sensory processing and motion sickness. While stereoacuity is predictive of behavioral performance in virtual reality, there seems to be no relationship between stereoacuity itself and either sense of presence or sickness [35], a result we replicated here as well. By contrast, as a group, those that prematurely quit viewing video content in virtual reality have exhibited greater sensitivity to 3D motion cues [17], a result that served as a primary motivator for the current study. Finally, beyond considerations of visual sensitivity, some studies have suggested that violations of sensory expectations contribute to motion sickness as well (e.g., [36]).

While the aforementioned studies explore explanations of variation in motion sickness susceptibility based on cue conflict theory, alternative accounts exist. Postural instability theory posits that motion sickness is instead due to an inability to regulate postural sway [37]. A number of studies report postural precursors of motion sickness in movement magnitude preceding the onset of motion sickness symptoms in physical environments [38], video games [39], and virtual reality headsets [27]. Additionally, other measures of movement that are orthogonal to magnitude have also been implicated in motion sickness such as the width of the multifractal spectrum [27] and temporal movement dynamics [40]. While the literature on the relationship between postural instability and motion sickness is extensive, there is some debate on the exact nature of the relationship. Some studies suggest postural predictors of motion sickness exist in movements recorded before participants are exposed to motion stimuli of any kind (e.g., [26]-[27], [41]-[47]). However, other studies suggest instead that changes in sway occur at the same time as motion sickness onset [8], [48]-[49].

A second claim made by advocates of the postural instability theory is that postural sway is inherently different in females than males. Consequently, the theory predicts that females should exhibit a greater propensity for motion sickness. Indeed, sex-based differences in motion sickness are well established outside of VR (e.g., 50). Prior work in VR has found evidence for such sex differences in motion sickness susceptibility as well [17], [27], but see [27],[51], which suggest that sex differences may be task-specific. We did not find a relationship between sex and motion sickness in our study. We speculate that sex differences in motion sickness may be modulated by additional factors. The current results suggest that inappropriate device calibration may exacerbate any inherent sex-based differences in susceptibility to motion sickness. Instead of relying on a default IPD and height as has been typical in prior research, we carefully calibrated the HMD to match the IPD and height of each observer.

The data presented here relied on the Oculus DK2 and one might ask to what extent motion sickness could be mitigated by newer generations of headsets with enhanced specifications (e.g., improved display properties, more comfortable designs, etc.). Our primary findings demonstrate that the extent to which one is sensitive to motion parallax information predicts one’s susceptibility to motion sickness. Thus, although the display quality and weight of the headset may certainly have elicited some discomfort, head-tracking and scene updating are expected to be more critical factors in general given our results. To our knowledge, the tracking specifications of newer devices are not dramatically different and, thus, the results presented here should generalize.

It has been suggested that administering a motion sickness questionnaire by itself creates an expectation that an observer will get sick, elevating scores upon repeated administration [52]. However, some observers reported little to no sickness after video viewing (see Figure 4), suggesting that the “inflation effects” of repeated SSQs reported by [52] were not shared by all participants in our study. Furthermore, when comparing sickness reports across all stages, the increase in sickness was not linear - responses after the 3D motion discrimination task exhibited a smaller increase on average relative to baseline, compared to the much more dramatic increase on average after video viewing both relative to the post-task and post-baseline reports, consistent with our a priori expectations. Finally, for the inflation effect to drive our results, the effect would need to have been larger for individuals with greater visual sensitivity to motion parallax cues, which we find difficult to justify. Therefore, while we cannot exclude that inflation might have occurred across repeated test administration in our design, it is unlikely that our results can be explained on that basis.

In conclusion, the current results account for variability in susceptibility to VR-induced motion sickness across individuals but imply that those individuals who would benefit the most from the visual cues that can be presented in VR are also those who may experience the most discomfort [17]. Our results suggest that motion sickness is not a “necessary evil” of VR technology. Our results motivate a number of strategies that can reduce sources of conflict and make the technology more accessible. First, VR experiences with modes that require less dramatic head movements by including alternative ways to complete tasks such as “teleporting” rather than navigating may offer more comfortable experiences. Second, experiences in which the intensity of the sickness-inducing cues is gradually increased over multiple exposures, can be an effective way to reduce motion sickness [53]-[54], thus slowly increasing the intensity of VR exposure may be an important recommendation for new users. Similarly, a somewhat counterintuitive option is to make the visual cues that induce motion sickness less reliable, by for example blurring, contrast reduction, or reducing the fidelity of the visual display through other means. Under such conditions observers will downweigh or even completely discount these cues, reducing the cue conflict signals produced by them, and therefore lower their susceptibility to motion sickness.

## Acknowledgements

We would like to thank Xuanxuan Ge and Elizabeth Shank for assistance with subject recruitment and data collection.

